# Structure of the ceramide-bound SPOTS complex

**DOI:** 10.1101/2023.02.03.526835

**Authors:** Jan-Hannes Schäfer, Carolin Körner, Bianca M. Esch, Sergej Limar, Kristian Parey, Stefan Walter, Dovile Januliene, Arne Moeller, Florian Fröhlich

## Abstract

Sphingolipids are structural membrane components that also function in cellular stress responses. The serine palmitoyl-transferase (SPT) catalyzes the rate limiting step in sphingolipid biogenesis. Its activity is tightly regulated through multiple binding partners, including Tsc3, Orm proteins, ceramides, and the phosphatidylinositol-4-phosphate (PI4P) phosphatase Sac1. The structural organization and regulatory mechanisms of this complex are not yet understood.

Here, we report the high-resolution cryo-EM structures of the yeast SPT in complex with Tsc3 and Orm1 (SPOT) as dimers and monomers and a monomeric complex further carrying Sac1 (SPOTS). In all complexes, the tight interaction of the downstream metabolite ceramide and Orm1 reveals the ceramide dependent inhibition. Additionally, observation of ceramide and ergosterol binding suggests a co-regulation of sphingolipid biogenesis and sterol metabolism within the SPOTS complex.

## Introduction

Sphingolipids are essential membrane components in eukaryotes, composed of a sphingosine backbone with a fatty acid attached, and are further modified through the addition of various polar head groups. They are particularly abundant in the plasma membrane, contributing to its structural integrity and function^1^. In addition, sphingolipids act as signaling molecules, for example, in apoptosis and the immune response^2–4^. Sphingolipid metabolism is tightly regulated via multiple signals and additionally linked to sterol levels^5–7^. Imbalances in the levels of sphingolipids and sterols are implicated in a variety of human pathologies, including neurodegenerative diseases such as Niemann-Pick type C and childhood amyotrophic lateral sclerosis (ALS)^8,9^.

Serine palmitoyl transferase (SPT) is the rate-limiting enzyme in the synthesis of sphingolipids. It catalyzes the transfer of a palmitoyl group to L-serine, yielding 3-ketosphinganine (3-KS; PMID: 3081509). 3-KS is reduced to long-chain bases, which are further processed into ceramides and complex sphingolipids^10^. SPT is highly conserved across species and consists of two large catalytic subunits (in yeast: Lcb1 and Lcb2) and interacts with a small regulatory subunit (in yeast: Tsc3)^11–16^. Tsc3 modulates enzyme activity through various mechanisms, including allosteric regulation and protein-protein interactions^17^.

Two recent structural studies of the human SPT showed that the enzyme acts as a homodimer with the two transmembrane helices of the human Lcb1 (SPTLC1) subunits swapped between dimers^18,19^. The small regulatory subunit ssSPTa provides an additional transmembrane helix, and the ORMDL3 protein is located in between the transmembrane helices of SPTLC1 and ssSPTa. The structures revealed the mechanism of substrate recognition and fatty acid selectivity. Regulation of the SPT complex occurs through multiple input signals, including Orm proteins, which are co-purified with the SPT^20^.

Orm proteins (ORMDL1/2/3 in mammalian cells, Orm1/2 in yeast cells) act as negative regulators of the SPT^20,21^. Mammalian ORMDL3 extends its N-terminus into the active site of SPT, thus inhibiting enzyme activity^18,19^. Yeast Orm proteins have extended N-termini that are not evolutionarily conserved, suggesting a different mechanism of regulation. In line, yeast Orm proteins are phosphorylated at the extended N-terminus by the Ypk kinases, leading to increased SPT activity^22–24^. In addition, SPT activity is also reduced in the presence of ceramides, the downstream metabolites of long-chain base/sphingosine synthesis. This regulation is proposed to depend on the presence of Orm proteins, but the molecular mechanism remains elusive^20,21,25^.

In yeast, the SPT-Orm-Tsc3 complex (SPOT) harbors an additional partner, the PI4P phosphatase Sac1 (SPOTS complex)^26–28^. Sac1 has been proposed to modulate the sphingolipid metabolism through its interaction with the SPOT complex; however, neither its binding mode nor its specific function within the SPOTS complex are known, but its deletion affects sphingolipid levels^29^.

Here, we solved cryo-EM structures of the yeast SPOT complex in both monomeric and dimeric states and the monomeric SPOTS complex. The overall architecture of the individual subunits is almost indistinguishable from yeast to human. A marked difference is the absence of the previously reported crossover helices at the protomer interface in our dimeric structure, which could explain why we were able to also obtain monomeric SPOT complexes. Notably, the PI4P phosphatase Sac1 only binds to the monomeric complex. Our data show that in yeast, Orm1 does not regulate SPT via insertion of its N-terminus in the active site but rather in conjunction with ceramide. We identified ceramide in all complexes, coordinated between Orm1 and Lcb1, blocking the SPT substrate channel. Furthermore, we revealed the presence of several ergosterol molecules in the monomeric complexes, suggesting that the SPOTS complex is a regulatory junction to coordinate sphingolipid and sterol levels. Together, we provide a structural basis for SPT regulation via multiple signals.

## Results

To unravel the architecture of the yeast SPOTS complex, we generated an *S. cerevisiae* strain co-expressing Lcb1, Lcb2, Tsc3, Orm1, and Sac1 under the control of the inducible GAL1 promotor. Lcb1 was internally FLAG-tagged after P9 to enable affinity purification while maintaining the functionality of Lcb1^13^. In addition, the three known phosphorylation sites S51, S52, and S53 of Orm1 were mutated to alanine to yield a non-phosphorylatable version (ORM1^AAA^). We reasoned that this would stabilize the entire complex. These three serine residues are target sites for the regulatory yeast Ypk kinase, which upon phosphorylation, increases SPT activity^20,22^. We anticipated that the ORM1^AAA^ mutant would render the SPOTS complex inactive; however, it still showed enzymatic activity of 45 nmol mg^−1^ min^−1^ and was sensitive to myriocin (sup. Fig. 1e).

For cryo-EM studies, the *S. cerevisiae* SPOTS complex was solubilized in glyco-diosgenin (GDN) and purified by FLAG-based affinity chromatography (sup. Fig. 1b). The purified complex was also subjected to mass spectrometric analysis, confirming the presence of all subunits with high sequence coverage (sup. Fig. 1d). Multi-model single particle cryo-EM revealed three different compositions of the complex within one dataset, including a C2 symmetric SPOT dimer (Fig. 1a) and two SPOT monomers, among which one additionally contains the regulatory subunit Sac1 (SPOTS complex) (Fig. 1b-c).

**Figure 1:**
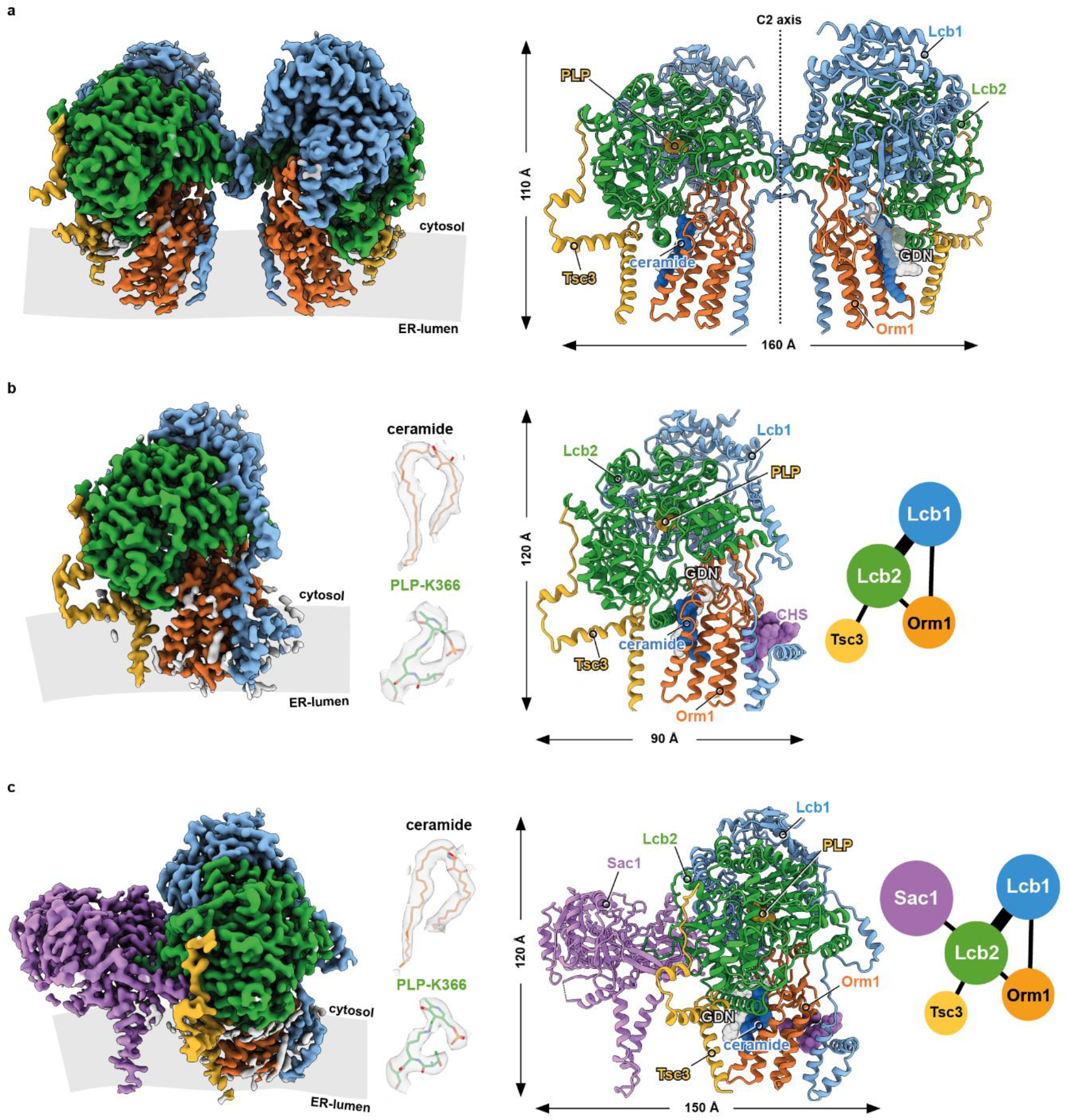
Overall architecture of yeast SPT complexes. **a** Cryo-EM density map and model of the C2 symmetric yeast SPOT dimer at 3.4 Å resolution. **b** The map and the model of the monomeric SPOT complex at 3.4 Å resolution. The middle inset shows the local densities for 44:0;4 ceramide, and PLP-K366. The contact diagram of the subunits within the complex is displayed on the right. The thickness of each line in the contact diagrams is proportional to the contact area between subunits. **c** The monomeric SPOTS complex at 3.3 Å resolution. The contact diagram is shown on the right, and the local densities for 44:0;4 ceramide, and PLP-K366 are in the middle. All individual subunits are consistently colored: Lcb1 in blue, Lcb2 in green, Orm1 in orange, Tsc3 in yellow, and Sac1 in purple. The membrane plane is indicated in gray.

### The architecture of the SPOT dimer

The SPOT dimer was refined to 3.4 Å resolution, with C2-symmetry imposed. Symmetry expansion of one protomer improved the resolution further to 3.0 Å (Tab.1 and Sup. Fig. 3,4). As previously reported, Lcb1 and Lcb2 build the enzymatic core of the complex. In contrast to the previously suggested architecture of yeast Lcb1^12^, only a single transmembrane helix (TM1) located at the N-terminal part of the protein is visible and anchors Lcb1 to the membrane, while Lcb2 is embedded in the membrane via an amphipathic helix (Fig. 2a, sup. Fig. 6, sup. Fig. 8c). The regulatory subunit Tsc3 provides an additional membrane anchor through its single transmembrane helix and an amphipathic helix (Fig. 2a, sup. Fig. 6). Tsc3 does not interact with Lcb1, but it binds tightly to Lcb2 via an elongated N-terminal region that is not conserved in the human homolog (Fig. 2d). Orm1 is positioned between the amphipathic helix of Lcb2 (Fig. 2g) and the TM1 helix of Lcb1 (Fig. 2i) but does not interact with Tsc3.

**Figure 2:**
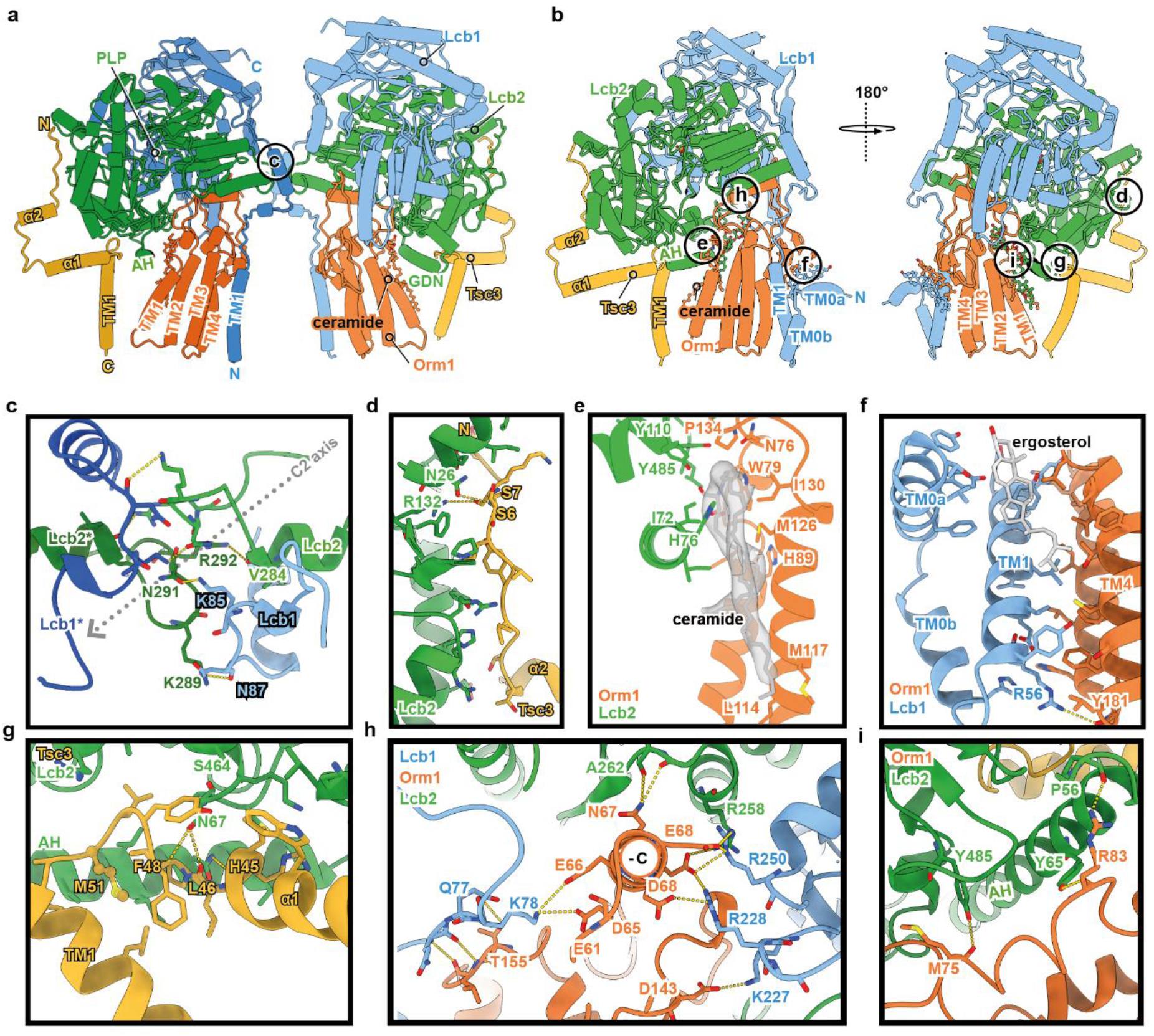
Overview of the specific subunit interactions within the SPOT complex. In the dimeric (a) and the monomeric (b) complexes, Orm1 interacts with Lcb1 and Lcb2. Lcb1 and Lcb2 interact tightly with each other, while Tsc3 binds exclusively to Lcb2. Colour code is the same as in Fig.1. The Lcb1 subunit of the second protomer is shown in a darker blue to provide a better view of the dimer interface. The circles indicate the areas of zoom-in views on the specific interactions in panels c-i. **c** Close-up view on the dimer interface, formed by the interactions between the cytosolic residues of Lcb1 and Lcb2. **d,g** Close-up views on the interactions between Tsc3 and Lcb2, **e** Ceramide 44:0;4 binding to Lcb2 and Orm1, **i** Interactions of the amphipathic helix AH of Lcb2 with Orm1, **h** Orm1 interactions with Lcb1 and Lcb2 via a short α-helix (-C = neg. electrostatic potential), **f** Ergosterol binding to Lcb1 and Orm1. Residues, which mediate key interactions, are shown as sticks. Polar contacts are indicated with yellow dotted lines. The subunits are depicted as cartoons, and ligands are shown in ball-and-stick representation. The experimental density for ceramide is shown in transparent gray.

The general architecture of the SPOT complex is remarkably conserved from yeast to human. However, our structure shows a different arrangement of the two Lcb1 transmembrane helices (TM1), which were previously reported to establish a crossover between the two protomers^18,19^, leading to an extensive interface within the membrane (sup. Fig. 7a). We do not observe such helix crossover for the yeast SPOT dimer. Nevertheless, the relative position of the Lcb1 transmembrane helix of one protomer superimposes with the corresponding crossover helix of the adjacent protomer in the human dimeric structure (sup. Fig. 7b-c). In yeast SPOT, the protomer binding interface is established through salt bridges between Lcb1^K85^ - Lcb2^N291^ and Lcb1^N87^ - Lcb2^K289^ and H-bonds between the adjacent Lcb2^V284^ - Lcb2*^R292^ (Fig. 2b) in the cytosol-facing portion of the protein. The distance between the Lcb1 transmembrane helices of our dimer structure is increased from 13 to 28 Å (sup. Fig. 7a), reminiscent of the previously reported ORMDL3-free SPT complex (PDB: 7K0I, 2.8 Å global RMSD).

The small human ssSPTa subunit has been shown to regulate fatty acid selectivity via the insertion of a methionine side chain in the substrate binding tunnel. The corresponding Tsc3 subunit in yeast harbors a methionine at a similar position between its transmembrane- and amphipathic helix, but its side chain does not extend into the substrate access channel (Fig. 2e). Interestingly, Tsc3 has been reported to regulate the amino acid choice of the SPT rather than controlling fatty acid selectivity as shown for ssSPTa^17^.

Superposition of ORMDL3 and Orm1 reveals very high structural conservation; only the regulatory N-terminal parts exhibit marked differences (Fig. 3c, sup. Fig. 9c). In the human SPT complex, the N-terminal methionine of ORMDL3 reaches into the substrate binding tunnel in SPTLC2, resulting in negative modulation of SPT activity. This requires a sharp kink from residue V10 to N11. The corresponding asparagine N73 in yeast is preceded by P72, which introduces a sharp kink in the opposite direction and folds into a structured helix, docking tightly into a socket composed of Lcb2 (Fig. 2h, 3c). Of note, the yeast Orm proteins harbor additional 63 N-terminal amino acids that are not present in the human proteins (sup. Fig. 9b-c). In our structure, the N-terminus of Orm1 latches onto the surface of Lcb1 before it interacts with its own C-terminus in an antiparallel beta-sheet (Fig. 3c). Consequently, the N-terminus cannot fulfill the same function that has been reported for the human SPT-ORMDL3, explaining the need for different modes of regulation that have been observed.

**Figure 3:**
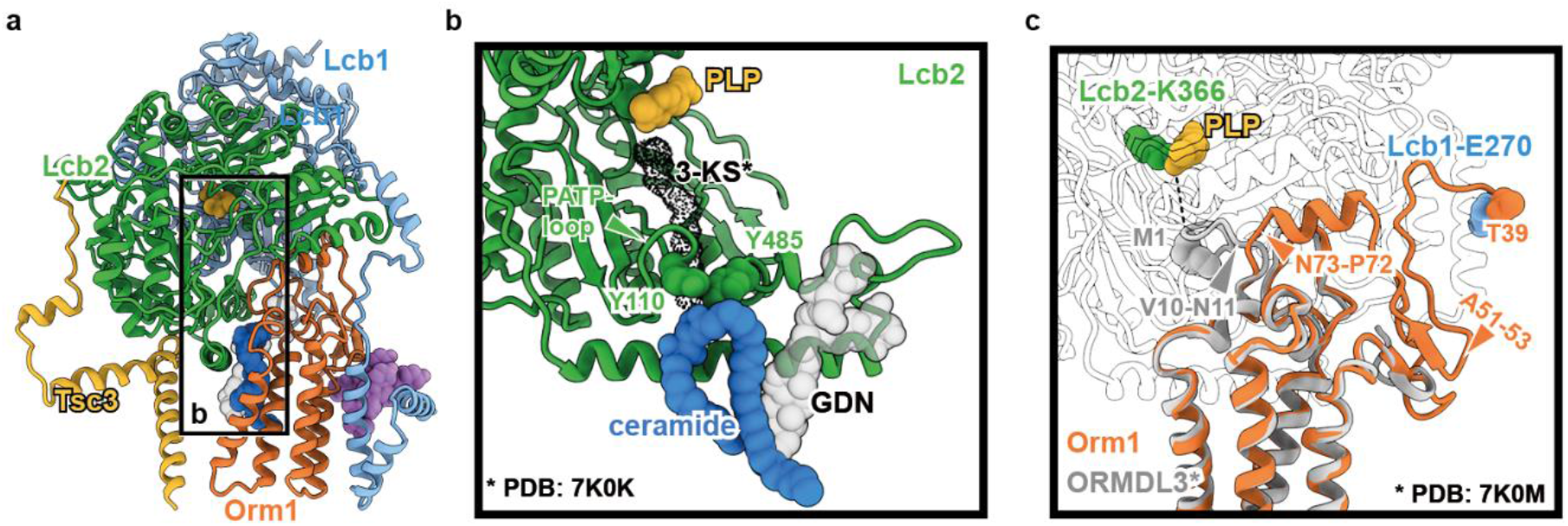
Regulation of SPT activity by Orm1 and ceramides. **a** Ligand binding within the SPOT complex. Colour code is the same as in Fig. 1. Ligands are represented as spheres with ergosterol in violet, 44:0;4 ceramide in dark blue, PLP in yellow, and GDN in semi-transparent gray. **b** Blocking of the substrate access tunnel by the putative Lcb2-gatekeeper residues Y110 and Y485 and ceramide 44:0;4, which is further stabilized by GDN. A docked 3-KS molecule (PDB: 7K0K, dotted black density) indicates the upper region of the substrate access tunnel. **c** Superposition of Orm1 and ORMDL3 (PDB: 7K0M) highlights divergence of M1-ORMDL3 towards the active site at Lcb2-K366 and Orm1-T39 towards Lcb1-E270. Diverging residues of ORMDL3 (V10-N11) and Orm1 (N73-P72) are marked with a triangle. Phosphorylation sites are indicated with an orange triangle (residues mutated from serine to alanine). Other subunits were omitted for clarity

As shown for the SPT-ORMDL3 structures, also in our structure, the active site between Lcb1 and Lcb2 is populated by the cofactor pyridoxal 5′-phosphate (PLP) (Fig. 1b-c, sup. Fig. 5b), which forms an internal aldimine with Lcb2-K366, essential for the catalysis of serine and acetyl-CoA condensation reaction^30^. Likewise, the putative substrate access tunnel is gated by the conserved PATP loop of Lcb2/SPTLC2 (in yeast, amino acids 486-489, Fig. 3b). Below the PATP loop, at the interface between the amphipathic helix of Lcb2^58-85^ and Orm1, we identified an elongated density (Fig.1b-c, Fig. 3b, sup. Fig. 4e), that was not detected in the human structures. Two long acyl chains and the lack of a prominent head group indicate the presence of a very long-chain fatty acid containing ceramide. To identify the bound ligand, the sample used for cryo-EM was subjected to lipid extraction and targeted lipidomics. As a control, an Orm-free preparation of the SPT-Tsc3-Sac1 complex was used. This analysis revealed the typical yeast 44:0;4 ceramide enriched in the SPOTS preparation, supporting its presence in the purified complex (sup. Fig. 1g). Ceramides, the downstream metabolites of the SPT catalyzed reaction, have been reported to negatively regulate SPOTS activity^25^. Therefore, the positioning of the 44:0;4 ceramide headgroup in the immediate proximity of the conserved, substrate-tunnel-gating PATP loop and direct interactions with Y485 and Y110 in Lcb2 (Fig. 2f) effectively blocks access to the substrate tunnel from the membrane, highlighting a regulatory mechanism of ceramide-based inhibition of the SPOT complex. In addition to 44:0;4 ceramide, we see a clear density for a GDN molecule, interacting with the long acyl chains of the ceramide via non-polar contacts (sup. Fig. 5a,d).

### The structure of the SPOT monomer

Previous studies were confined to dimeric SPT-complex preparations, and monomeric structures arise from focused classification and refinements. The crossover helices between the SPTLC1 subunits in the SPT-ORMDL3 structures offer a simple explanation of why the purification of individual monomeric complexes has not been possible previously.

Superposition of the monomeric SPOT complex, solved to 3.4 Å resolution (Fig. 1b), with the dimeric version reveals only minor overall differences (RMSD = 0.48 Å across all 554 pairs, sup. Fig. 7f). Most notably is an interrupted transmembrane helix at the N-terminus of Lcb1, that is absent in humans (sup. Fig 9d). In humans, the N-terminus of SPTLC1 starts with a short amphipathic helix located in the ER lumen, consecutively entering the membrane as the previously mentioned crossover helix. While the position of the corresponding transmembrane helix Lcb1-TM1 is virtually identical in yeast, the organization of the N-terminus is very different (Fig. 2c). Preceding the TM1, Lcb1 folds into a short transmembrane helix (TM0b) that spans approximately half of the bilayer (15 Å in length) and leads through a short loop into another helix, that runs parallel to the membrane and is deeply embedded in the upper leaflet (TM0a). This places the Lcb1 N-terminus in the cytosol, which was confirmed by its ability to recruit a cytosolic GFP-tagged ALFA nanobody to the ER membrane (suppl. Fig 10). The TM0a helix also forms a hydrophobic pocket with the Orm1-TM4, in which three structurally well-resolved ergosterol molecules are positioned (Fig. 2i, Sup. Fig. 5c). The N-terminal helices are not resolved in the dimeric SPOT complex, and superposition of two SPOT-monomer models onto each C2 symmetric protomer within the SPOT dimer model results in a sterical clash of adjacent Lcb1-TM0a (sup. Fig. 7f). The presence of sterol molecules was confirmed by an enzyme-coupled reaction, resulting in the detection of ~25 ng ergosterol per μg protein from the purified complex (sup. Fig. 1f). The direct interaction of the SPT with ergosterol offers a potential explanation for the co-regulation of sterols and sphingolipids in yeast, as discussed before^6^.

### Structure of the SPOTS complex

Previous works have identified the interaction of the PI4P phosphatase Sac1 with the SPOT complex; however, the role of Sac1 within the complex remains elusive. Here, we solved the structure of the SPOTS complex at 3.3 Å resolution (Fig. 1c). In the crystal structure of the Sac1 phosphatase domain, large parts of this domain have not been resolved and thus interpreted as flexible regions (PDB: 3LWT)^28^. In our cryo-EM structure, Sac1 tightly interacts with the flexible N-terminal loop of Lcb2 and is anchored to the lipid bilayer via two transmembrane helices and an amphipathic helix (Fig. 4a, sup. Fig. 6). Membrane-embedded Sac1 within the SPOTS complex does not show significant flexibility. Sac1 interacts with Lcb2 exclusively, and we do not observe a previously suggested interaction with Tsc3^31^. The binding interface between Sac1 and Lcb2 involves the eight amino-terminal residues of Lcb2 and a C-terminal hairpin β-sheet motif in Sac1 (Fig. 4c, sup. Fig. 9a). A smaller binding interface involves a H-bond between Sac1^I104^ - Lcb2^N88^ and several non-polar contacts (Fig. 4b). Interestingly, Sac1 is only bound to monomeric SPOT complexes in our data.

**Figure 4:**
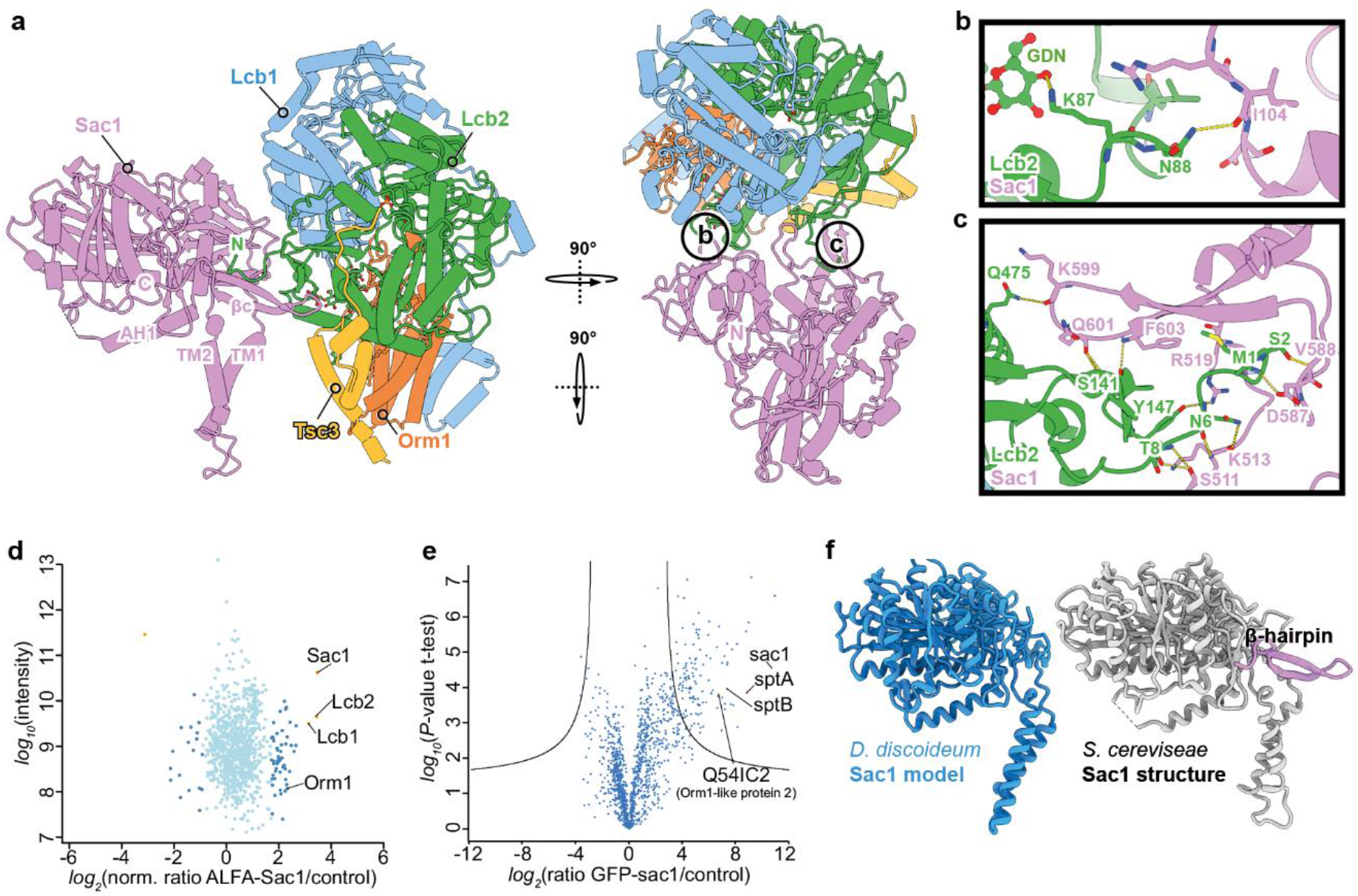
Sac1 binding to the SPOT complex. **a** Sac1 interactions within the SPOTS complex. Colour code is the same as in Fig. 1. **b-c**, Close-up views of the interactions between Sac1 and Lcb2. Polar contacts are indicated with yellow dotted lines. The subunits are depicted as cartoons, and ligands are shown in ball-and-stick representation. **d** Proteomic analysis of ALFA-Sac1 compared to untagged control cells. Protein intensities are plotted against heavy/light SILAC ratios. Significant outliers are colored in red (p<1^−11^), orange (p<1^−4^), or steel blue (p<0.05); other proteins are shown in light blue. **e** Label-free proteomics of *D. discoideum* cells expressing GFP-Sac1 compared to untagged control cells. In the volcano plot, the protein abundance ratios of GFP-Sac1 over control cells are plotted against the negative log_10_ of the *P*-value of the t-test for each protein. **f** Comparison of AlphaFold prediction of *D. discoideum* Sac1 (Q55AW9) with the yeast Sac1 from our cryo-EM structure. Yeast-specific β-hairpin motif is highlighted in violet.

To investigate local sequence conservation within the Sac1-Lcb2-interface, homologs from human and *D. discoideum*, lacking the C-terminal hairpin β-sheet motif, were used for multiple sequence alignment, which revealed poor conservation of key-residues across species (sup. Fig. 7e, 9a). However, a superposition of the AlphaFold model of *D. discoideum* Sac1 with our experimental structure suggests a conserved basis for Lcb2-binding independent of the Sac1 β-sheet motif (sup. Fig. 7d).

To test if the SPOTS complex also exists in other organisms, we compared the interaction partners from ALFA-Sac1 pulldowns in yeast (Fig 4d) with GFP-Sac1 pulldowns from *D. discoideum* (Fig 4e) using mass spectrometry-based proteomics. These experiments revealed the presence of the SPOTS complex in both organisms. *S. cerevisiae* Sac1 showed interactions with Lcb1, Lcb2, and the Orm1 protein. *D. discoideum* Sac1 showed interactions with the two SPT subunits, sptA and sptB, and the Orm1-like protein 2. The β-hairpin motif of *S. cerevisiae* Sac1 can thus be interpreted as a yeast-specific regulatory structure.

## Discussion

The SPT enzyme is the rate-limiting factor in sphingolipid metabolism and controls cellular sphingolipid homeostasis through multiple input signals. Here, we present the structure of the yeast SPOT complex as both a monomer and a dimer and also the SPOTS complex, including the PI4P phosphatase Sac1. The SPOT complex is highly conserved across different species, as evidenced by the similarities in structure between yeast and mammalian SPT-Orm complexes. However, we also observe marked differences that explain their different regulatory mechanisms.

The lack of the Lcb1-TM1 helix swap, previously observed in human SPT complexes, results in weaker protomer interactions and explains the presence of monomeric complexes. Both oligomeric states from the human complexes are active *in vitro*; however, their respective physiological relevance is unclear. Importantly, we only detected the regulatory subunit Sac1 in interaction with the monomer. Co-purification of Orm1 and Orm2 is low when the complexes are purified through the Orm subunits^20^. It is also difficult to conceive that the cell is able to discriminate between the two highly homologous Orm proteins during the loading of a dimeric SPOT complex. Finally, Orm2 is exclusively regulated through the endosome/Golgi-associated degradation (EGAD) pathway^32^. Together, this suggests that the monomeric SPOT and SPOTS complexes are the predominant forms in yeast.

The membrane-spanning helices of Lcb1 in our structures differ largely from the currently annotated membrane topology. Biochemical studies suggested the presence of three transmembrane helices, which are spread across far-apart regions of Lcb1 (amino acids 50-84, 342-371, 425-457)^12^. In all of our structures, the yeast Lcb1 TM1 is located at the same position as the S1 helix of human SPTLC1. Additionally, we also find one more interrupted N-terminal membrane inserted helix (TM0a: T20-Q35 and TM0b: Q40-S49) in front of the TM1 helix in the monomeric species.

Despite the high structural resemblance, the Orm-dependent SPT regulation is different in yeast and humans. In human cells, the N-terminus of ORMDL3 regulates access to the substrate binding pocket of SPTLC2^18^. In yeast, both Orm1 termini face away from the active site of Lcb2, requiring another mechanism of regulation. Additionally, the phosphorylation of three serine residues on the N-terminal loop of Orm1/2 has been discussed to influence SPT activity^22–24^. Notably, these amino acids are located at a highly accessible loop, connecting the N-terminus of Orm1 to Lcb1. Therefore, it can be anticipated that phosphorylation at this position leads to the rearrangement of the complex, potentially affecting its stability or activity.

Furthermore, our data explain the ceramide-induced regulation that has been reported for yeast SPT complexes, which is dependent on Orm binding^25^. In all of our structures, TM3-4 of Orm1 act as a docking station for a 44:0;4 ceramide, which is sandwiched between Orm1 and the amphipathic helix of Lcb2. The ceramide extends out of the cytosol-facing membrane plane through interactions with two aromatic residues, Y110 and Y485, of Lcb2. We propose that the two tyrosine residues function as gatekeepers controlling access to the previously discovered substrate channel^18^. The Orm1 protein would thus inflict its inhibitory function only in conjunction with ceramide. We also observe a decrease in ceramide levels in Orm1-free SPT samples, further supporting the Orm1 mediated regulatory effect of ceramide.

It is an interesting observation that ceramide is not the only co-purified lipid in the monomeric SPOT and SPOTS complexes. The deeply embedded helix Lcb1-TM0a directly interacts with three molecules of ergosterol in a position that would correspond to the upper leaflet of the ER membrane. Levels of ergosterol and sphingolipids have been previously suggested to be tightly connected to each other^6,7,33^. It is appealing to speculate that SPT activity could be directly regulated via the levels of ER-localized ergosterol. Since the superposition of two monomeric structures onto a dimeric complex causes steric clashes of TM0a helices (sup. Fig. 7f), it is possible that high levels of ergosterol promote the formation of the TM0a helix, breaking the dimer apart and thus changing SPT activity.

Sac1 has been shown to interact with the SPOT complex and was proposed to affect its activity^20^. The canonical function of Sac1 is the dephosphorylation of PI4P in the ER. PI4P is generated at the Golgi apparatus and is exchanged with ER-synthesized sterols via the oxysterol binding proteins (OSBPs)^34–37^. This exchange is driven by the Sac1-dependent dephosphorylation of PI4P in the ER. We show that Sac1 specifically interacts with the N-terminus of the Lcb2 subunit and not with Tsc3, as suggested previously^31^. A short C-terminal β-hairpin structure supports the interaction but might also have a yeast-specific regulatory function since it is not present in other species. The structure of Sac1 includes two transmembrane helices and an additional amphipathic helix. This shows that the previously uncharacterized region between the phosphatase domain and the transmembrane helices is structured when in contact with a hydrophobic moiety, proving that Sac1 can only act at the membrane where it is inserted^28,38–41^.

In summary, we present a detailed picture of the protein interactions within the SPOTS complex, which is at the heart of neurological disorders such as HSAN1 and childhood ALS^9,42–44^. We reveal how SPT activity is controlled through its downstream metabolite ceramide. The additional ergosterol binding site provides the first mechanistic link in the co-regulation of sphingolipid and sterol metabolism.

## Material and Methods

### Yeast strains

Yeast strains and plasmids used in this study are listed in supplementary tables 1 and 3. For purifications, SPOTS subunits were expressed under the control of the GAL1 promoter using integrative plasmids. The 3x-FLAG tag was inserted between codons 9 and 10 of LCB1, as previously reported^20^.

### Purification of 3xFLAG tagged SPOTS complex from *S. cerevisiae*

Yeast cells were collected after growth for 24 h in yeast peptone (YP) medium containing 2 % galactose (v/v), washed in lysis buffer (50 mM HEPES-KOH (pH 6.8), 150 mM potassium acetate (KOAc), 2 mM MgOAc, 1 mM CaCl_2_, 200 mM sorbitol) and resuspended in a 1:1 ratio (w/v) in lysis buffer supplemented with 1 mM phenylmethylsulfonylfluoride (PMSF) and 1x FY protease inhibitor mix (Serva). Resuspended cells were frozen in a drop-by-drop fashion in liquid nitrogen, pulverized in 15x 2 min cycles at 12 CPS in a 6875D Freezer/Mill Dual-Chamber Cryogenic Grinder (SPEX SamplePrep), and thawed in lysis buffer with 1 mM PMSF and 1x FY. After two centrifugation steps at 1,000 *g* at 4 °C for 20 min, microsomal membranes were pelleted at 44,000 *g* at 4 °C for 30 min. Cells were resuspended in lysis buffer and then diluted with IP buffer (50 mM HEPES-KOH, pH 6.8, 150 mM KOAc, 2 mM MgOAc, 1 mM CaCl_2_, 15 % Glycerol) with 1% glyco-diosgenin (GDN) supplemented with protease inhibitors. After nutating for 1.5 h at 4 °C, unsolubilized membranes were pelleted at 44,000 *g* at 4 °C for 30 min. The supernatant was added to α-Flag resin (Sigma Aldrich) and nutated for 45 min at 4 °C. Beads were washed twice with 20 ml IP buffer with 0.1 % GDN and 0.01 % GDN, respectively. Bound proteins were eluted twice on a turning wheel in IP buffer with 0.01 % GDN for 45 min and 5 min, respectively, at 4 °C with 3xFLAG peptide. The eluates were collected by centrifugation at 1,800 rpm, at 4 °C for 30 s and concentrated in a 100 kDa Amicon Ultra centrifugal filter (Merck Millipore), which was equilibrated with 1% GDN in IP buffer. The concentrated eluate was applied to a Superose 6 Increase 5/150 column (Cytiva) for size exclusion chromatography (SEC) and eluted in 50 μl fractions using ÄKTA go purification system (Cytiva). Peak fractions were collected, concentrated as described before, and used for further analysis.

### Cryo-EM sample preparation and data acquisition

Sample quality was inspected by negative-stain electron microscopy as previously described^45^. Micrographs of the negatively-stained sample were recorded manually on a JEM2100plus transmission electron microscope (Jeol), operating at 200 kV and equipped with a Xarosa CMOS (Emsis) camera at a nominal magnification of 30,000, corresponding to a pixel size of 3.12 Å per pixel.

For cryo-EM, the sample was concentrated to 10 mg/ml. C-flat grids (Protochips; CF-1.2/1.3-3Cu-50) were glow-discharged, using a PELCO easiGlow device at 15 mA for 45 s and 3 μl of the concentrated sample were immediately applied and plunge frozen in liquid ethane, using a Vitrobot Mark IV (Thermo Fisher) at 100 % relative humidity, 4 °C. The dataset was collected using a Glacios microscope (Thermo Fisher), operating at 200 kV and equipped with a Selectris energy filter (Thermo Fisher) with a slit of 10 eV. Movies were recorded with a Falcon 4 direct electron detector (Thermo Fisher) at a nominal magnification of 130,000 corresponding to a calibrated pixel size of 0.924 Å per pixel, and the data was saved in the electron-event representation (EER) format. The dose rate was set to 5.22 e^−^ per pixel per second and a total dose of 50 e^−^ per Å^2^. 13,604 movies were collected automatically, using EPU software (v.2.9, Thermo Fisher) with a defocus range of −0.8 to −2.0 μm.

### Cryo-EM image processing

The SPOTS dataset was processed in cryoSPARC (v.4), and the processing workflow is presented in supplementary Fig. 2. Movies were preprocessed with patch-based motion correction, patch-based CTF estimation and filtered by the CTF fit estimates using a cutoff at 5 Å in cryoSPARC live (v.4), resulting in a remaining stack of 12,552 micrographs.

Well-defined 2D classes were selected and used for subsequent rounds of template-based 2D classification. 1.006.434 particles were extracted in a box of 432 pixels, and Fourier cropped to 216 pixels. Previously generated ab initio 3D reconstructions from live processing were used for several rounds of heterogeneous refinement, which resulted in two distinct, well-defined reconstructions. Both reconstructions were processed separately.

#### SPOT-dimer-complex

Classes corresponding to the newly termed SPOT-dimer-complex were subjected to non-uniform refinement, and 3D-aligned particles were re-extracted without binning. Additional rounds of heterogeneous refinement following non-uniform refinement and local refinement with C2 symmetry applied resulted in a stack of 142K particles and a consensus map with 3.4 Å resolution.

To further improve the map quality, particles were symmetry expanded by using a C2 point group, thereby aligning signals coming from both protomers onto the same reference map. Subsequent signal subtraction and local refinement results were subjected to 3D classification in PCA mode focused around the protomer. Additional local CTF correction and local refinement of 94,884 symmetry-expanded particles yielded a focused map with an overall resolution of 3.0 Å.

#### SPOTS-monomer

Similarly, a stack of 252,688 particles was re-extracted without binning using the alignment shifts from a heterogeneous refinement. Aligned particles were further classified through heterogeneous refinement and 3D classification in PCA mode. A final round of non-uniform-refinement with higher-order aberration correction enabled, local CTF correction, and local refinement resulted in an overall resolution of 3.3 Å.

#### SPOT-monomer

An additional well-resolved 3D class was identified, lacking the Sac1-assigned density, and processed separately. 123K particles were cleaned through 2D classification to remove the remaining non-protomer classes. A stack of 96K particles was subjected to heterogeneous refinement, and the remaining 89K particles were further refined through nonuniform and local refinement. This resulted in a final map with a global resolution of 3.4 Å.

All maps were subjected to unsupervised B-factor sharpening within cryoSPARC. Reported B-factors resulted from un-supervised auto-sharpening during refinement in cryoSPARC. To aid model building, unsharpened half-maps were subjected to density modification within Phenix *phenix.resolve_cryo_em*.

### Model building and refinement

Initial atomic models were generated using the AlphaFold2 prediction of a monomer, which was placed in the individual maps and fitted as rigid bodies through UCSF Chimera^46^. The structure was manually inspected in Coot (v.0.9)^47^ and iteratively refined using *phenix.real_space_refine* within Phenix (v.1.19). Validation reports were automatically generated by MolProbity^48^ within Phenix^49^. All density maps and models have been deposited in the Electron Microscopy Data Bank and the PDB. The PDB IDs are 8C82 (SPOTS-Dimer-Complex), 8C80 (SPOTS-Orm1-Monomer) and 8C81 (SPOTS-Orm1-Monomer-Sac1). The respective EMDB IDs are EMD-16469, EMD-16467, and EMD-16468. All structural data was visualized with ChimeraX^46^ and protein interactions were analyzed with the help of PDBe PISA^50^. 2D ligand-protein interaction diagrams were calculated in LigPlot^+ 51^. Characterization of selected helices was performed in HeliQuest^52^.

### Proteomics

Proteomics analysis was performed as described previously^53^. Briefly, cells were grown in SDC medium containing either light or heavy lysine (30 μg/ml)^54^. Main cultures were inoculated in 500 ml SDC medium containing light or heavy lysine (30 μg/ml) from a pre-culture to an OD_600_ and grown over day to an OD_600_ 0.8. The same amount of OD units from both cultures was harvested at 4,000 rpm, at 4 °C for 5 min, resuspended in ALFA pull-down buffer (20 mM HEPES pH 7.4, 150 mM potassium acetate, 5% glycerol, 1x FY and 1 mM PMSF) and frozen in a drop-by-drop fashion in liquid nitrogen. Cells were lysed with 500 μl acid-washed glass beads in 500 μl ALFA pull-down buffer using the FastPrep (MP biomedicals). After a centrifugation step at 1,000 *g* and 4 °C for 10 min, GDN was added to a final concentration of 1 %, and the supernatants of light and heavy lysine cultured strains were incubated for 1.5 h at 4 °C on a turning wheel. Supernatants were spun down at 14,000 rpm at 4 °C for 10 min and incubated on a turning wheel at 4 °C with 12.5 μl in ALFA pull-down buffer equilibrated ALFA beads (NanoTag Biotechnologies). The beads were washed in total six times, first with pull-down buffer and then with washing buffer (20 mM HEPES pH 7.4, 150 mM potassium acetate, 5% glycerol). Beads of light and heavy lysine cultured strains were combined during the last washing step. Proteins on the beads were reduced, alkylated, and digested with LysC at 37 °C overnight following the protocol of the iST Sample Preparation Kit (PreOmics) for protein digestion. Dried peptides were resuspended in 10 μl LC-Load, and 5 μl were used to perform reversed-phase chromatography on a Thermo Ultimate 3000 RSLCnano system connected to a QExactive*PLUS* mass spectrometer (Thermo Fisher Scientific) through a nano-electrospray ion source. Peptides were separated on a PepMap RSLC C18 easy spray column (2 μm, 100 Å, 75 μm x 50 cm, Thermo Fisher Scientific) with an inner diameter of 75 μm and a column temperature of 40 °C. Elution of peptides from the column was realized via a linear gradient of acetonitrile from 12-35% in 0.1% formic acid for 80 min at a constant flow rate of 250 nl/min following a 20 min increase to 60%, and finally, 10 min to reach 90% buffer B. Eluted peptides were directly electro sprayed into the mass spectrometer. Mass spectra were acquired on the Q Exactive*Plus* in a data-dependent mode to automatically switch between full scan MS and up to ten data-dependent MS/MS scans. The maximum injection time for full scans was 50 ms, with a target value of 3,000,000 at a resolution of 70,000 at *m/z* 200. The ten most intense multiply charged ions (z≥2) from the survey scan were selected with an isolation width of 1.6 Th and fragmented with higher energy collision dissociation^55^ with normalized collision energies of 27. Target values for MS/MS were set at 100,000 with a maximum injection time of 80 ms at a resolution of 17,500 at m/z 200. To avoid repetitive sequencing, the dynamic exclusion of sequenced peptides was set at 20 s. Resulting data were analyzed using MaxQuant (version 2.2.0.0, www.maxquant.org/)^56,57^, the R software package (www.r-project.org/; RRID:SCR_001905), and Perseus (V2.0.7.0, www.maxquant.org/perseus)^58^. The mass spectrometry proteomics data have been deposited to the ProteomeXchange Consortium via the PRIDE^59^ partner repository. Data will be made available upon request.

### Ceramide analysis of purified SPOTS complex

Ceramide was extracted and measured from 20 μg of purified proteins. 150 mM ammonium formate was added to the purified complex. As an internal standard, ceramide (CER d17:1/24:0; Avanti) was added, and lipid extraction with 2:1 chloroform/methanol was performed as described previously^60,61^. Dried lipid films were dissolved in a 50:50 (v/v) mixture of Buffer A (50:50 water/acetonitrile, 10 mM ammonium formate, and 0.1% formic acid) and B (88:10:2 2-propanol/acetonitrile/water, 2 mM ammonium formate and 0.02% formic acid). An external standard curve was prepared using phytoceramide (CER t18:0/24:0; Cayman Chemical). Samples were analyzed on an Accucore C30 LC column (150 mm x 2.1 mm 2.6 μm Solid Core; Thermo Fisher Scientific) connected to a Shimadzu Nexera HPLC system and a QTRAP 5500 LC-MS/MS (SCIEX) mass spectrometer. Different lipid species (CER d17:1/24:0 as a control (Avanti Polar lipids), CER 42:0;3, CER 42:0;4, CER 42:0;5, CER 44:0;3, CER 44:0;4, CER 44:0;5 and PC 16:0/18:1) were detected in a positive MRM mode with optimized transition settings within a 6 min HPLC run. For peak integration, the SciexOS software was used. The concentrations of all ceramide species were calculated using the external standard curve. The ceramide concentrations were expressed in pmol/μg protein.

### GFP-Trap pull-down from *Dictyostelium discoideum*

*Dictyostelium discoideum* strains (sup. Tab. 2) expressing either pDM317-GFP-Sac1 or pDM317-GFP^62^ were grown at 22°C in HL5-C medium (ForMedium) supplemented with geneticin (G418, 5 μg/ml). Electroporation of *D. discoideum* was performed according to Paschke et al. 2018 with modification^63^. The cell number was determined (Countess II F2, Thermo Fisher Invitrogen), and 3 ∙ 10^7^ cells were used for each sample. Pull-downs were performed in triplicates. The cells were pelleted and washed once in cold Soerensen-Sorbitol.

Cell pellets were snap frozen in Eppendorf tubes and lysed with glass beads in 500 μl GFP pull-down buffer (20mM HEPES pH 7.4, 150mM KOAc, 5% glycerol, 1% GDN, Roche Complete Protease Inhibitor Cocktail EDTA free, Roche) using a FastPrep (MP biomedicals). The supernatant was cleared at 14,000 rpm for 10 min and incubated for 10 min rotating at 4 °C together with 12.5 μl pre-equilibrated GFP-Trap beads (Chromotek). Beads were washed four times with GFP pull-down buffer at 2,500 *g* for 2 min at 4 °C. Afterwards, they were washed two times with wash buffer (20mM HEPES pH 7.4, 150mM KOAc, 5% glycerol) at 2,500 *g* for 2 min at 4 °C. Beads were further treated following the “iST Sample Preparation Kit (Agarose Immunoprecipitation Samples)” protocol with the iST Sample Preparation Kit (PreOmics) for protein digestion. Dried peptides were resuspended in 10 μl LC-Load, and 4 μl were used to perform reversed-phase chromatography as described above. The mass spectrometry proteomics data have been deposited to the ProteomeXchange Consortium via the PRIDE^59^ partner repository. Data will be made available upon request.

### Colorimetric-based enzymatic assay

SPT activity was measured by monitoring the release of CoA-SH from the SPT-catalyzed condensation of palmitoyl-CoA and L-serine. All assays were performed on a 200-μl scale. Briefly, all assays were performed in IP buffer with 0.008 % GDN, 15 mM L-serine, 100 μM palmitoyl-CoA, and 30 μM PLP. The reaction was initiated by adding 1 μg protein. To validate SPT activity, 100 μM myriocin was added to inhibit SPT activity. Since myriocin was dissolved in methanol (MeOH), an appropriate amount of MeOH was added to all other samples. After incubation at RT for 1 h, the samples were deproteinized with a 3 kDa MWCO concentrator (Merck Millipore). To measure CoA levels, the samples were further treated in a 96-well plate following the “Coenzyme A Assay Kit” protocol from Sigma-Aldrich. The absorbance of the colorimetric product (570 nm) was measured in a SpectraMax iD3 Multi-Mode microplate reader. Corrected absorbances were obtained by subtracting the absorbance of the protein-free samples from the absorbances of the protein-containing samples.

### Fluorescence-based ergosterol measurements from purified protein

Bound ergosterol levels were measured using an enzymatic coupled assay generating the highly fluorescent dye resorufin. All assays were performed on a 100-μl scale in a 96-well plate according to the “Amplex™ Red Cholesterol Assay Kit” protocol (ThermoFisher Scientific). No cholesterol esterase was added. Fluorescence was recorded (λ_EX_=550nm, λ_EM_ =590nm) after three hours using the SpectraMax iD3 Multi-Mode microplate reader. Relative fluorescence intensity was obtained by subtracting the fluorescence intensity of the protein-free samples from the intensity of the protein-containing samples.

### Fluorescence microscopy

Cells were grown overnight at 30 °C in synthetic medium supplemented with essential amino acids (SDC), diluted in the morning to an OD_600_ of 0.15, and grown to logarithmic growth phase. Cells were directly imaged live in SDC medium using a Zeiss Axioscope 5 FL (Zeiss) equipped with a Plan-Apochromat 100x (1.4 numerical aperture (NA)) oil immersion objective and an Axiocam 702 mono camera. Data were acquired with ZEN 3.1 pro software and processed with ImageJ 2.1.0. (National Institutes of Health, Bethesda, MD; RRID:SCR_003070). Single medial planes of yeast cells are shown.

**Table 1:** Cryo-EM data collection, refinement and validation statistics.

|  | SPOT-Dimer | SPOT-dimer<br>masked | SPOTS | SPOT-monomer |
| --- | --- | --- | --- | --- |
| **Data Collection** |  |  |  |  |
| Accession number | EMD-16469 | EMD-16485 | EMD-16468 | EMD-16467 |
| Magnification | 130,000 | 130,000 | 130,000 | 130,000 |
| Voltage / kV | 200 | 200 | 200 | 200 |
| Dose / e-Å<sup>-2</sup> | 50 | 50 | 50 | 50 |
| Pixel size / Å | 0.924 | 0.924 | 0.924 | 0.924 |
| Defocus range / µm | -2.0 to -0.8 | -2.0 to -0.8 | -2.0 to -0.8 | -2.0 to -0.8 |
| Recorded movies | 13,604 | 13,604 | 13,604 | 13,604 |
| Final particle images | 141,900 | 94,884 | 53,236 | 89,484 |
| Microscope | Glacios | Glacios | Glacios | Glacios |
| Camera | Falcon 4 | Falcon 4 | Falcon 4 | Falcon 4 |
| Energy Filter | Selectris | Selectris | Selectris | Selectris |
| **Image Processing** |  |  |  |  |
| Initial model | AlphaFold 2 | AlphaFold 2 | AlphaFold 2 | AlphaFold 2 |
| Processing software | cryoSPARC (v.4) | cryoSPARC (v.4) | cryoSPARC (v.4) | cryoSPARC (v.4) |
| Symmetry imposed | C2 | C2 | C1 | C1 |
| Resolution (FSC0.143) / Å | 3.4 | 3.0 | 3.3 | 3.4 |
| Applied B-factor / Å<sup>2</sup> | -105 | -86 | -63 | -79 |
| **Model Refinement** |  |  |  |  |
| PDB accession | 8C82 |  | 8C81 | 8C80 |
| Validation |  |  |  |  |
| FSCmap-to-model(0.143) / Å | 3.0 |  | 3.2 | 3.3 |
| MolProbity score | 1.68 |  | 1.45 | 1.33 |
| Clash Score | 7.93 |  | 3.98 | 3.71 |
| Composition |  |  |  |  |
| Atoms | 21,208 |  | 15,938 | 10,968 |
| Protein residues | 2,634 |  | 1,966 | 1,350 |
| Ligands | 6 |  | 6 | 6 |
| Bonds (R.M.S.D.) |  |  |  |  |
| Length (Å) | 0.003 |  | 0.003 | 0.003 |
| Angles (°) | 0.653 |  | 0.541 | 0.615 |
| B-factors (min/max/mean) |  |  |  |  |
| Protein residues | 50.62/143.88/63.21 |  | 60.65/221.62/100.09 | 54.22/176.06/87.84 |
| Ligand | 23.35/152.04/114.63 |  | 83.24/117.15/105.19 | 90.15/120.51/106.88 |
| Ramachandran plot (%) |  |  |  |  |
| Favored | 96.33 |  | 96.06 | 97.02 |
| Allowed | 3.67 |  | 3.94 | 2.98 |
| Outliers | 0.00 |  | 0.00 | 0.00 |
| Rotamer outliers (%) | 0.00 |  | 0.00 | 0.00 |

## Acknowledgments

We thank Jürgen Heinisch (Osnabrück) for providing the GFP-tagged ALFA-nanobody plasmid. We thank Oliver Schmidt (Innsbruck) for providing the original 3xFLAG-Lcb1 plasmid. We thank Caroline Barisch (Osnabrück) for sharing her expertise in *Dictyostelium discoideum* cell culture and the Hilbi laboratory (Zürich) for the pDM317 plasmids expressing GFP-Sac1 and GFP. This work was funded by the DFG (SFB944 P20 to FF and P27 to AM, the SFB1557 P6 to FF, P11 to AM, FR 3647/2-2 and FR 3647/4-1 to FF, INST.190/196-1 FUGG), BMBF 01ED2010 to AM. JHS is supported by a fellowship from the Friedrich Ebert foundation.

## Author contributions

Investigation: JHS, CK, SL, BE, SW. Formal analysis: JHS, CK, SL, BE, SW, KP Visualization: JHS, CK, KP, FF. Conceptualization: FF, AM, DJ. Funding acquisition: FF, AM. Writing—original draft: JHS, CK, FF, AM, DJ. Writing—review and editing: JHS, CK, FF, AM, DJ.

