## Supplementary-Information for "Structure of the ceramide-bound SPOTS complex"

[1] Osnabrück University
Department of Biology/Chemistry
Structural Biology section
49076 Osnabrück, Germany

[2] Osnabrück University
Department of Biology/Chemistry
Bioanalytical Chemistry section
49076 Osnabrück, Germany

[3] Osnabrück University
Center of Cellular Nanoanalytic Osnabrück (CellNanOs)
49076 Osnabrück, Germany

[†] These authors contributed equally to this work

[#] For correspondence:
 (F. F.); (A. M.);
 (D. J.)

**Keywords:** SPOTS complex, Sac1, serine palmitoyl-transferase, sphingolipids, ceramide

### Supplementary Figures and Tables

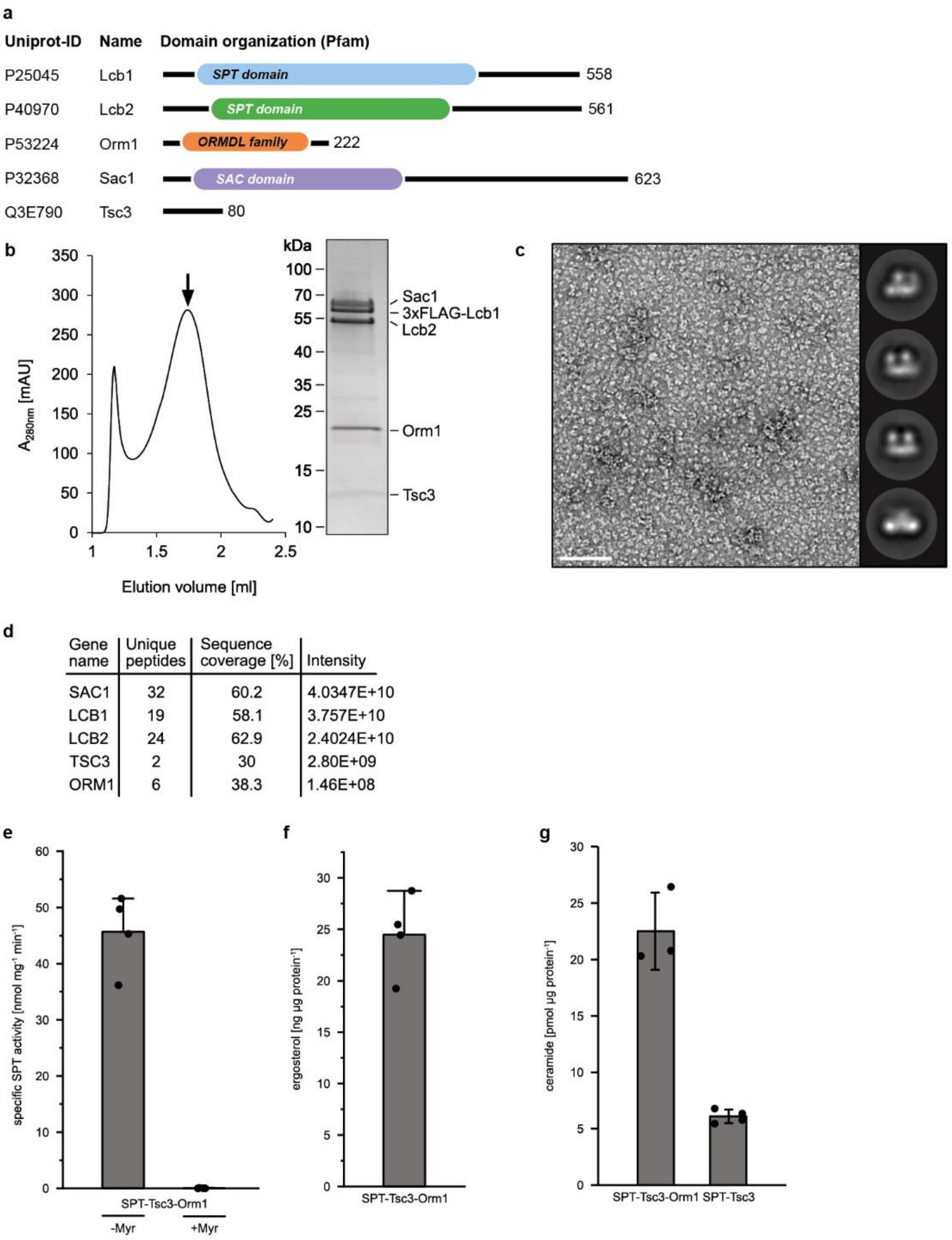


| Gene name | Unique peptides | Sequence coverage [%] | Intensity |
|---|---|---|---|
| SAC1 | 32 | 60.2 | 4.0347E+10 |
| LCB1 | 19 | 58.1 | 3.757E+10 |
| LCB2 | 24 | 62.9 | 2.4024E+10 |
| TSC3 | 2 | 30 | 2.80E+09 |
| ORM1 | 6 | 38.3 | 1.46E+08 |

Sup-Fig 1: ***In vitro* functional characterization of the yeast SPOTS complex**

**a** Pfam-based domain annotation of the SPOTS subunits. **b** SEC profile and corresponding Coomassie-blue stained SDS-PAGE gel of indicated fraction of GDN solubilized SPOTS complex. **c** Representative micrograph and 2D class averages from negative-stain TEM. 100 nm scale bar. **d** Summary of the mass-spectrometric analysis of the sample in **b**. **e** Specific SPT enzyme activity measurement with or without the specific SPT-inhibitor myriocin (Myr; four technical replicates). **f** Ergosterol quantification from GDN-solubilized and purified SPOT complex. **g** Ceramide quantification from GDN-solubilized and purified SPOT complex and Orm-free SPT-Tsc3.

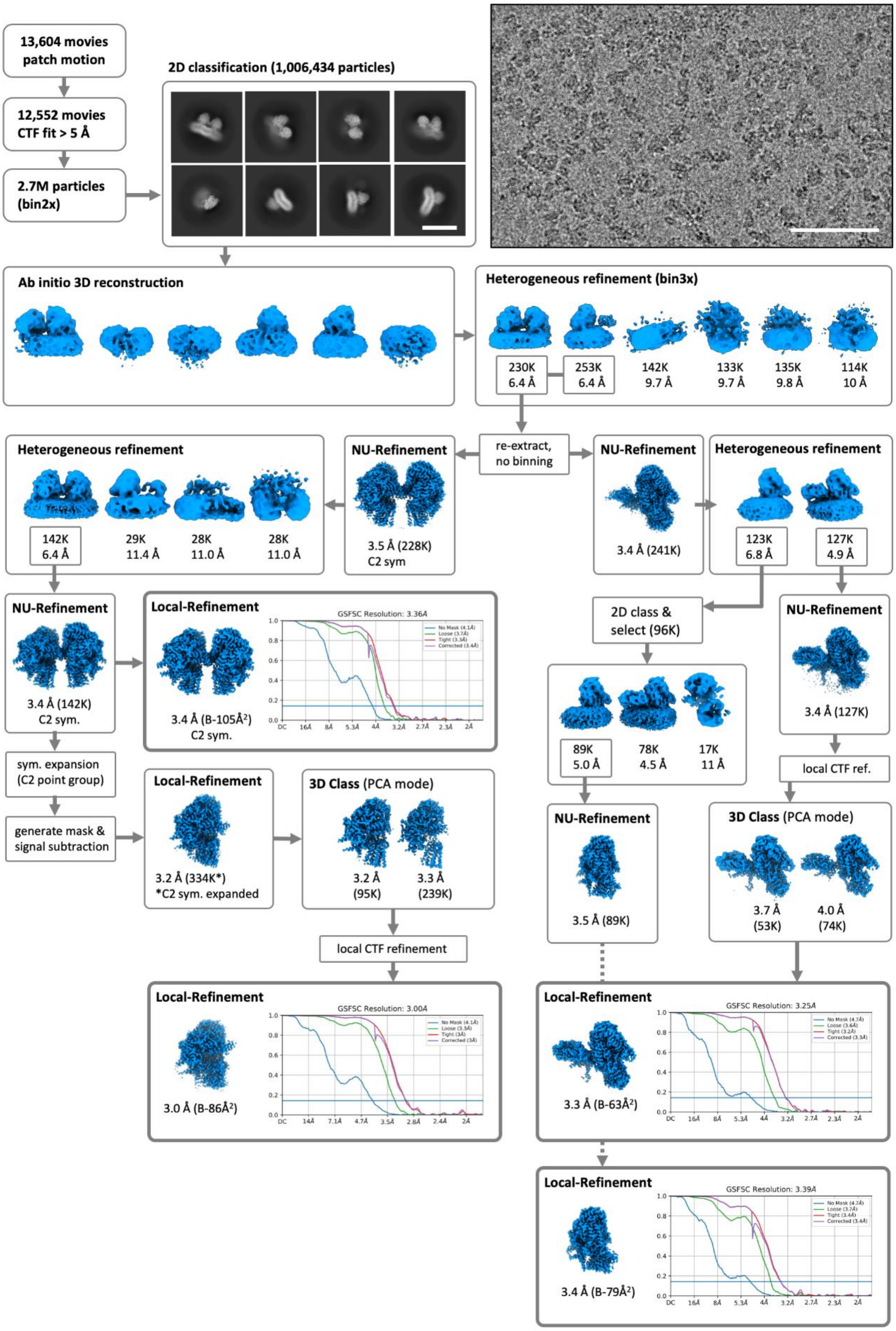
13,604 movies
patch motion
12,552 movies
CTF fit > 5 Å
2.7M particles
(bin2x)
2D classification (1,006,434 particles)
Ab initio 3D reconstruction
Heterogeneous refinement (bin3x)
230K 6.4 Å
253K 6.4 Å
142K 9.7 Å
133K 9.7 Å
135K 9.8 Å
114K 10 Å
re-extract, no binning
Heterogeneous refinement
142K 6.4 Å
29K 11.4 Å
28K 11.0 Å
28K 11.0 Å
NU-Refinement
3.5 Å (228K) C2 sym
NU-Refinement
3.4 Å (241K)
Heterogeneous refinement
123K 6.8 Å
127K 4.9 Å
NU-Refinement
3.4 Å (142K) C2 sym.
Local-Refinement
3.4 Å (B-105Å²) C2 sym.
GSFSC Resolution: 3.36Å
2D class & select (96K)
89K 5.0 Å
78K 4.5 Å
17K 11 Å
NU-Refinement
3.4 Å (127K)
local CTF ref.
sym. expansion (C2 point group)
generate mask & signal subtraction
Local-Refinement
3.2 Å (334K*)
*C2 sym. expanded
3D Class (PCA mode)
3.2 Å (95K)
3.3 Å (239K)
NU-Refinement
3.5 Å (89K)
3D Class (PCA mode)
3.7 Å (53K)
4.0 Å (74K)
local CTF refinement
Local-Refinement
GSFSC Resolution: 3.00Å
3.0 Å (B-86Å²)
Local-Refinement
GSFSC Resolution: 3.25Å
3.3 Å (B-63Å²)
Local-Refinement
GSFSC Resolution: 3.39Å
3.4 Å (B-79Å²)

Sup-Fig 2: **Cryo-EM analysis of the SPT-Orm1 complexes**

Processing workflow for SPOT-dimer, SPOT-monomer and SPOTS complex. All processing steps were performed in cryoSPARC. Representative cryo-EM micrograph and 2D-class averages. 100 nm scale bar in micrograph and 20 nm scale bar in 2D class-averages.

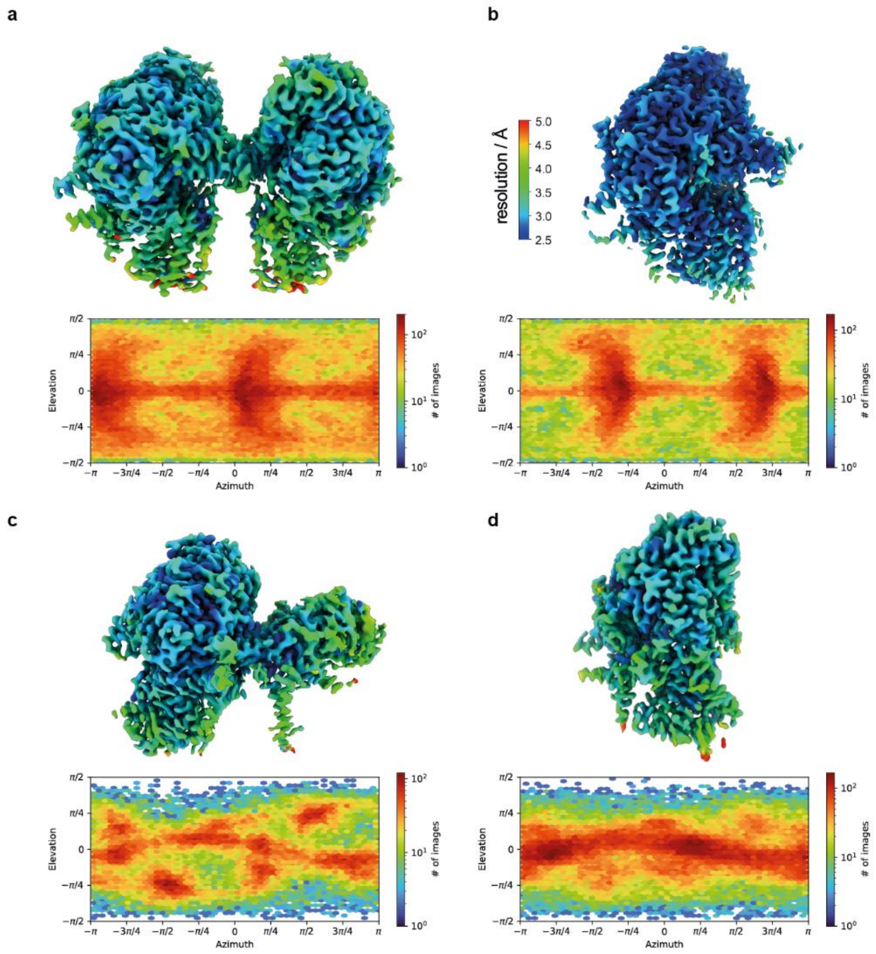


Sup-Fig 3: **Cryo-EM map evaluation**

Local resolution estimation and angular particle distribution of **a** SPOT-dimer, **b** masked SPOT-dimer, **c** SPOTS and **d** SPOT-monomer.

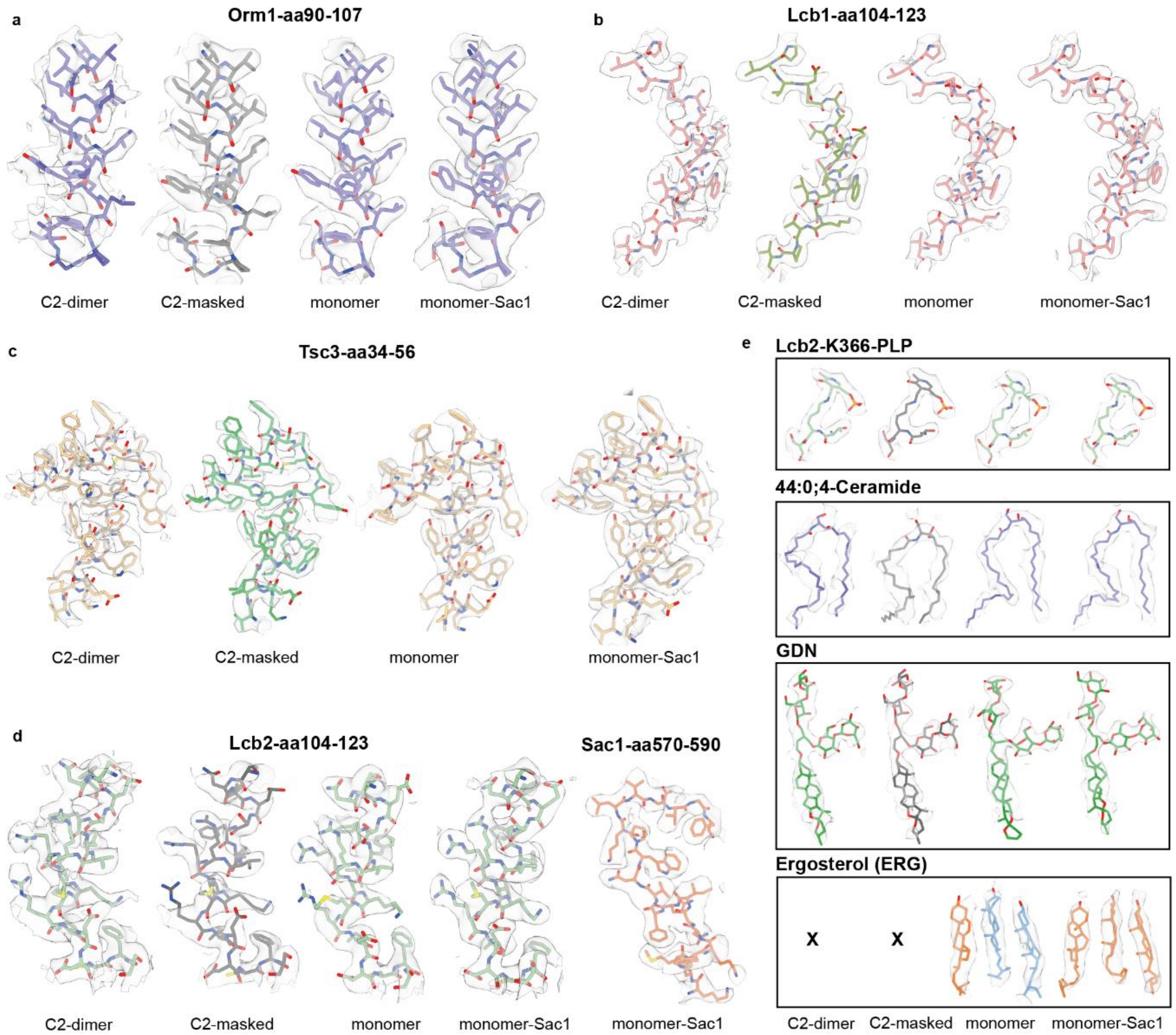


Sup-Fig 4: **Local cryo-EM density map quality of SPOT and SPOTS complexes**

Side-by-side comparison of selected residues and ligands within all cryo-EM density maps of **a** Orm1, **b** Lcb1, **c** Tsc3, **d** Lcb2 , Sac1 and ligands in **e**. Ergosterol (ERG) was not present in maps denoted with an x.

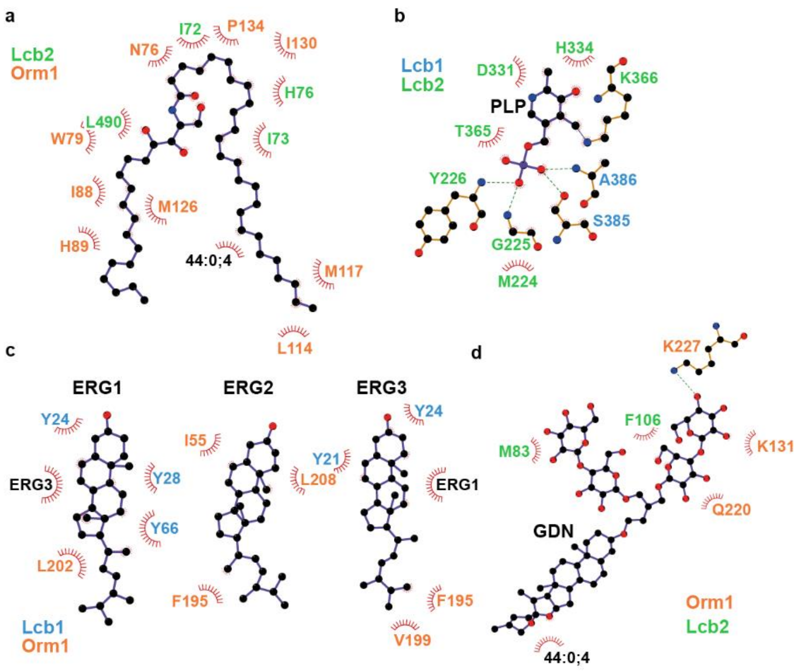


Sup-Fig 5: **Ligand interaction diagram of SPOT and SPOTS complexes**

2D ligand interaction diagram of ceramide 44:0;4 (**a**), internal aldimine between PLP and Lcb2$^{K366}$ (**b**), ergosterols (ERG) (**c**) and glyco-diosgenin (GDN) (**d**). Key interacting residues are shown as ball and sticks with polar contacts given as green dotted lines. Diagrams were calculated and visualized with LigPlot+. (a,b,d for all oligomers, c for SPOT-monomer and SPOTS).

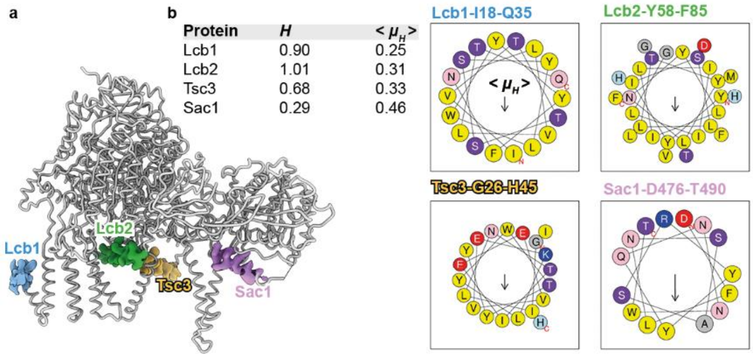


Sup-Fig 6: **Amphipathic helix (AH) analysis of yeast Lcb1, Lcb2, Tsc3 and Sac1**

**a** Identification of four amphipathic helices in SPOTS involves Lcb1$^{I18\text{-}Q35}$, Lcb2$^{Y58\text{-}F85}$, Tsc3$^{G26\text{-}H45}$ and Sac1$^{D476\text{-}T490}$. **b** Physico-chemical properties, including the mean hydrophobic moment $< \mu_H >$ (arrow proportional, perpendicular to membrane plane) and hydrophobicity $H$ were calculated with HeliQuest[52]. Helix wheels are colored according to their chemical polarity.

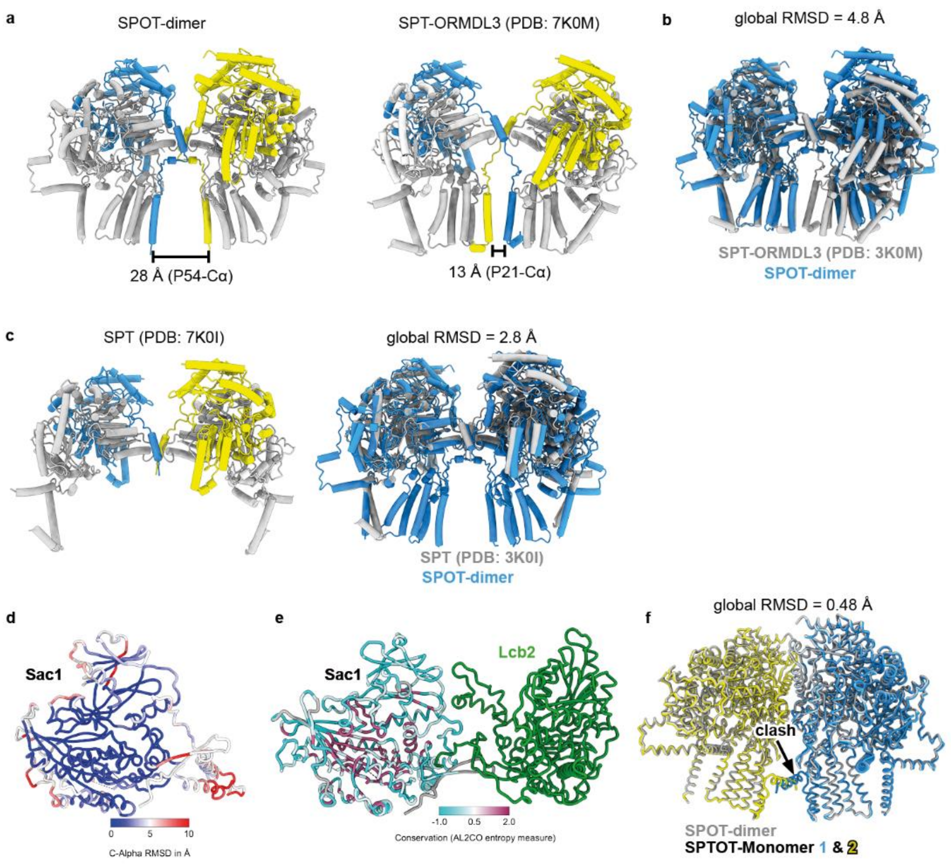


Sup-Fig 7: **Structural conservation within different SPOT oligomers**

**a** Side-by-side comparison of the Cα distances between the conserved proline residue in TM1 in the yeast SPOT-dimer and human SPT-ORMDL3 dimer (PDB: 7K0M). Adjacent subunits are colored in blue and yellow for clarity. **b** Superposition of SPT-ORMDL3 and SPOT-dimer results in a global root-mean-square deviation (RMSD) of 4.8 Å. **c** Overview and superposition of ORMDL3-free human SPT (PDB: 7K0I) and the SPOT-dimer results in a global RMSD of 2.8 Å. **d** Cα RMSD between the Sac1 subunit from SPOTS and an AlphaFold prediction of *Dictyostelium discoideum* (Q55AW9). **e** Sequence conservation between yeast Sac1 and its homolog from *D. discoideum*. The Lcb2 subunit (green) is added for highlighting the interface. **f** Superposition of two SPOT-monomers (blue and yellow) onto the SPOT-dimer (grey). The sterical clash of Lcb1-TM0a is indicated with an arrow.

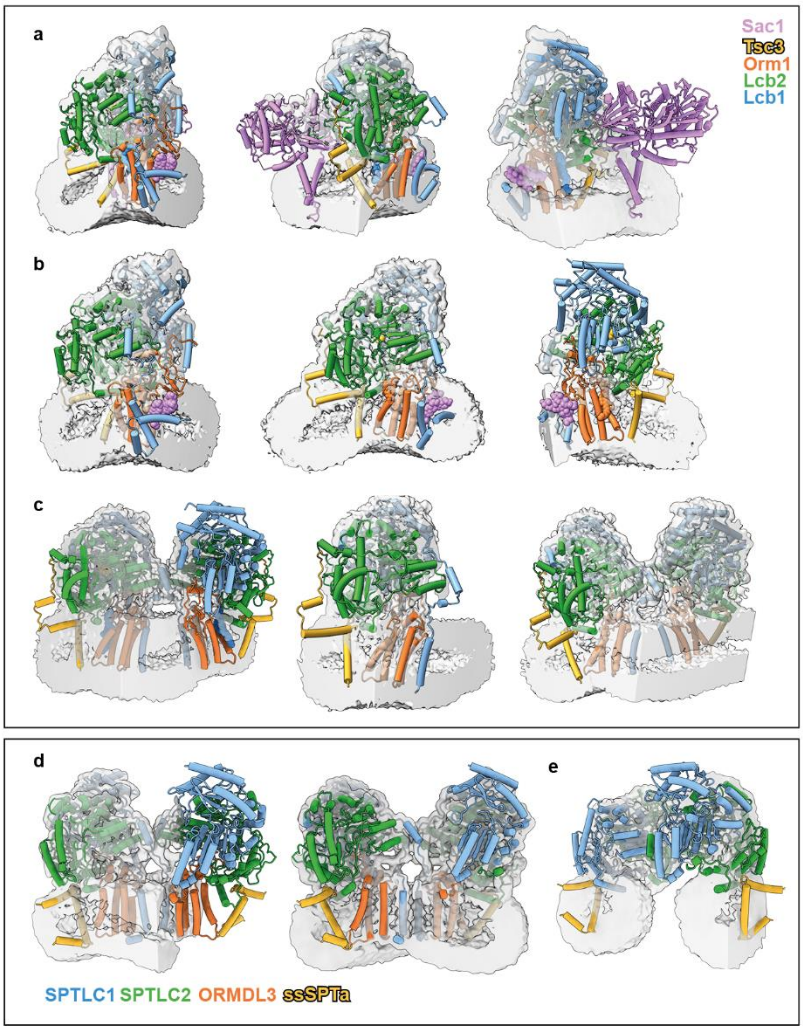


Sup-Fig 8: **Micelle curvature induction by SPT-ORMDL3 and yeast SPT oligomers**

Representative volume slices of **a** SPOTS **b** SPOT-monomer **c** SPOT-dimer **d** SPT-ORMDL3 (PDB: 7K0M, EMD-22602) and **e** SPTLC1/SPLTC2/ssSPTa (PDB: 7K0I, EMD-22598). EMDB maps were gaussian filtered by 1.5-fold standard deviations.

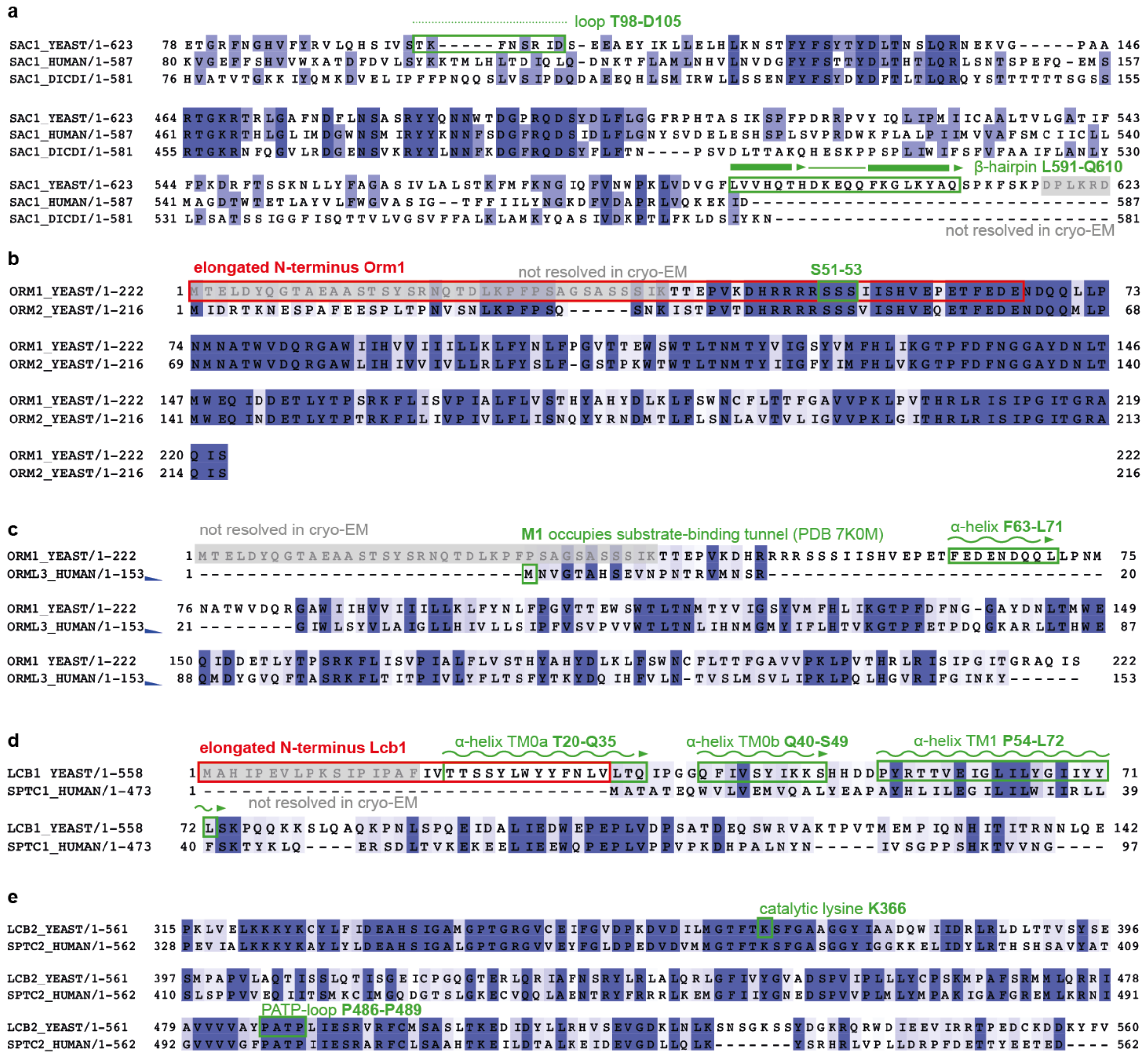


Sup-Fig 9: **Sequence conservation and unique features within different SPT complexes**

Clustal-based multiple-sequence alignment (MSA) of **a** Sac1 from *S. cerevisiae* (P32368) *H. sapiens* (Q9NTJ5) and *D. discoideum* (Q55AW9) with the C-terminal β-hairpin region and Lcb2 interacting loop T98-D105 highlighted in green. **b** Comparison of *S. cerevisiae* Orm1 (P53224) and Orm2 (Q06144). The elongated N-terminal region of Orm1 is colored in red and the mutated phosphorylation-site S51-53 is marked in green. **c** Comparison of *H. sapiens* ORMDL3 (Q53FV1) and yeast Orm1. The N-terminal methionine is highlighted in green, which was observed to occupy the substrate binding tunnel in human SPT-ORMDL3 (PDB: 7K0M). The Orm1 helix F63-L71 interacts with Lcb1/2 is highlighted in green. **d** Yeast Lcb1 (P40970) has an elongated N-terminal region, including helix TM0a (T20-Q35) and TM0b (Q40-S49), which are not present in human SPTLC1 (O15270). Non-resolvable residues from cryo-EM are marked in light-gray. Sequences are color coded by conservation with a cutoff at 30 %. MSAs were prepared with Jalview[63].

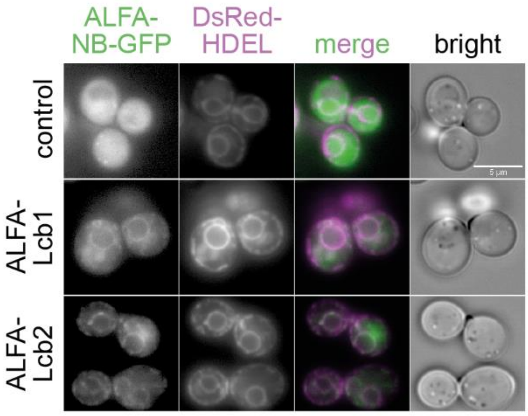


Sup-Fig. 10: **The N-terminus of Lcb1 faces the cytosol**

A cytosolic GFP-tagged ALFA nanobody (ALFA-NB-GFP) is expressed in control cells (upper panels), ALFA-Lcb1 cells (middle panels) and ALFA-Lcb2 cells (lower panels) also expressing DsRed-HDEL. Both, ALFA-Lcb1 and ALFA-Lcb2 allow the recruitment of the otherwise cytosolic ALFA-NB-GFP to the ER membrane marked with DsRed-HDEL showing that the N-termini of both proteins face the cytosol. GFP (left panels), DsRed-HDEL (middle left panels), merged images (middle right panels) and brightfield images (right panels) are shown. Scale bar = 5 µM.

Sup-Tab. 1: **List of all yeast strains used in this study**

| **Strain** | **Genotype** | **Reference** |
|---|---|---|
| FFY3085 | MATα *leu2-3,112 trp1-1 can1-100 ura3-1 ade2-1 his3-11,15 lcb1Δ::hphNT1 pRS415-3xFLAG-Lcb1::LEU2* | This study |
| FFY4354 | MATα *leu2-3,112 trp1-1 can1-100 ura3-1 ade2-1 his3-11,15 lcb1Δ::hphNT1 pRS415-3xFLAG-Lcb1::LEU2 sac1Δ1-264::kanMX6 pRS403-ALFA-Sac1::HIS3* | This study |
| FFY4192 | MATa/MATα *leu2-3,112 trp1-1 can1-100 ura3-1 ade2-1 his3-11,15 orm1Δ::natNT2 orm2Δ::hphNT1 pRS404-GAL1-Tsc3::TRP1 pRS406-GAL1-Orm1*$^{AAA}$*::URA3 pRS406-GAL1-3xFLAG-Lcb1::URA3 pRS405-GAL1-Lcb2::LEU2 pRS403-GAL1-Sac1::HIS3* | This study |
| FFY4133 | MATa/MATα *leu2-3,112 trp1-1 can1-100 ura3-1 ade2-1 his3-11,15 orm1Δ::natNT2 orm2Δ::hphNT1 pRS404-GAL1-Tsc3::TRP1 pRS406-GAL1-3xFLAG-Lcb1::URA3 pRS405-GAL1-Lcb2::LEU2 pRS403-GAL1-Sac1::HIS3* | This study |
| FFY5161 | MATα *leu2-3,112 ura3-52 his3-Δ200 trp1-Δ901 ADE2 suc2-Δ9 GAL lys2-801 lcb2Δ::hphNT1 pRS403-ALFA-Lcb2::HIS3 YIp128-ALFAnb-GFP::LEU2 pRS404-DsRed-HDEL::TRP1* | This study |
| FFY5162 | MATα *leu2-3,112 ura3-52 his3-Δ200 trp1-Δ901 ADE2 suc2-Δ9 GAL lys2-801 lcb1Δ::hphNT1 pRS406-ALFA-Lcb1::URA3 YIp128-ALFAnb-GFP::LEU2 pRS404-DsRed-HDEL::TRP1* | This study |
| FFY5228 | MATα *leu2-3,112 ura3-52 his3-Δ200 trp1-Δ901 ADE2 suc2-Δ9 GAL lys2-801 YIp128-ALFAnb-GFP::LEU2 pRS404-DsRed-HDEL::TRP1* | This study |

Sup-Tab. 2: **List of all *Dictyostelium discoideum* lines used in this study**

| **Strain** | **Plasmid used for transformation** | **Reference** |
|---|---|---|
| GFP-Sac1 | pDM317-GFP-Sac1 | Barisch lab (Osnabrück) |
| GFP | pDM317-GFP | Barisch lab (Osnabrück) |

Sup-Tab. 3: **List of all plasmids used in this study**

| Plasmid | Reference |
|---|---|
| pRS415-3xFLAG-Lcb1::LEU2 | Schmidt lab (Innsbruck) |
| pRS403-ALFA-Sac1::HIS3 | This study |
| pRS406-GAL1-Orm1$^{AAA}$::URA3 | This study |
| pRS404-GAL1-Tsc3::TRP1 | This study |
| pRS403-GAL1-Sac1::HIS3 | This study |
| pRS405-GAL1-Lcb2::LEU2 | This study |
| pRS406-GAL1-3xFLAG-Lcb1::URA3 | This study |
| pRS404-DsRed-HDEL::TRP1 | This study |
| YIp128-ALFAnb-GFP::LEU2 | Heinisch lab (Osnabrück) |
| pRS403-ALFA-Lcb2::HIS3 | This study |
| pRS406-ALFA-Lcb1::URA3 | This study |
| pDM317-GFP-Sac1 (G418) | Vormittag *et al.*, 2023 |
| pDM317-GFP (G418) | Vormittag *et al.*, 2023 |

Sup-Tab 4: **List of all oligo nucleotides used in this study**

| Name | Sequence |
|---|---|
| ALFA tag rev | TGG TTC GGT TAA TCT TCT TC |
| Sac1_aa2_for | ACA GGT CCA ATA GTG TAC |
| pRS406_for_Orm1 | ATGACCGAATTAGATTATCAAGGAACTG |
| pRS406_rev_Orm1 | GGCCCTACGCGCTCTAGA |
| GAL1_for_OH_Orm1promoter | CTAGAGCGCGTAGGGCCAGTACGGATTAGAAGCCGC |
| GAL1_rev_OH_ORM1 | TAATCTAATTCGGTCATGTTTTTTCTCCTTGACGTTAAAG |
| pRS404_GAL1_for | TATATCTAGAACTAGTGGATCCCC |
| pRS404_GAL1_rev | GTTTTTTCTCCTTGACGTTAAAG |
| TSC3_OH_GAL1pr_for | CGTCAAGGAGAAAAAACATGACACAACATAAAAGCTCG |
| TSC3_OH_pRS404_rev | GGGGATCCACTAGTTCTAGATATATCTGTGACTCGGATATGGAG |
| pRS403_Sac1_for | ATGACAGGTCCAATAGTGTAC |
| pRS403_Sac1pr_rev | CGATCAGGACGTCAGGG |
| GAL1_OH_Sac1pr_for | CCCTGACGTCCTGATCGAGTACGGATTAGAAGCCG |
| GAL1_OH_Sac1_rev | CACTATTGGACCTGTCATGTTTTTTCTCCTTGACGTTAAAG |
| pRS405_Lcb2_for | ATGAGTACTCCTGCAAACTATACC |
| pRS405_Lcb2_rev | CACCCAATCACCGCGCTT |
| GAL1pr_OH_Lcb2pr_for | AAGCGCGGTGATTGGGTGAGTACGGATTAGAAGCCG |
| GAL1pr_OH_Lcb2_rev | TTTGCAGGAGTACTCATGTTTTTTCTCCTTGACGTTAAAG |

| | |
|---|---|
| pRS406_FLAG_LCB1_for | ATGGCACACATCCCAGAG |
| pRS406_FLAG_LCB1_rev | GACAGAGCAGTATGTGAGG |
| GAL1pr_OH_Lcb1pr_for | CCTCACATACTGCTCTGTCAGTACGGATTAGAAGCCG |
| GAL1pr_OH_Lcb1pr_rev | TCTGGGATGTGTGCCATGTTTTTCTCCTTGACGTTAAAG |
| SAC1_KO_Rev | CAGCCCAGTATATTGGCACAGATCCTCTTGTCTGTAAGAAGGAGATCGATGAATTCGAGCTCG |
| SAC1_S1 new | ATAATATTTATATACACGTATATTTTCTCGTCTAGATATGcgtacgctgcaggtcgac |
| Lcb1_S1 | GTT ATT TAT CCT TTT TTC TTC CTT CCC ACC CAA AAA AAA AAA GCA ATG CGT ACG CTG CAG GTC GAC |
| LCB1 S2 | ATA TAT ATG TGC GTG TGC ATA TAC TGG CTT TCT ATT TTT AAT CGA TGA ATT CGA GCT CG |
| Orm1_S2 | AAA ATA TAA ATA TAG CAA AAA CAT CTA GAT ACA AGA TTG AAA TAA ACT ATG TTC AAT CGA TGA ATT CGA GCT CG |
| ORM1_S1 | AAG CAG AGT TAT TCT TAT TTT GTA TTT CAT TGC ATT TTT ATC CAT TTA GTT AAT GCG TAC GCT GCA GGT CGA C |
| ORM2_S1 | GAA TTA ACG CAA GAC TAT ACC ATT ATA AAA ACG CAT AAG AAA CAG TTT CAT CAT GCG TAC GCT GCA GGT CGA C |
| Orm2_S2 | Orm2 S2 |
| Lcb2_S1 | AAGATTCCACACACTTTATTGTGATAGTTTTCAAAGTAAAAAGTAATAGATTATGCGTACGCTGCAGGTCGAC |
| Lcb2_S2 | ACGTCTTCCAGAAATTTTGTAATTTTTCACCTAACTAGCAATTAGGTAAAATTCAATCGATGAATTCGAGCTCG |
| backbone_exchange_vec_for | GGCGTAATCATGGTCATAGC |
| backbone_exchange_vec_rev | CTCACTGGCCGTCGTTTTAC |
| backbone_exchange_ins_for | GTAAAACGACGGCCAGTGAG |
| backbone_exchange_ins_rev | GCTATGACCATGATTACGCC |
| pALFA-Lcb1-Q5-for | TTACGTCGTCGTTTGACCGAACCCAAATCAATACCGATTCCGG |
| pALFA-Lcb1-Q5-rev | TTCCTCTTCCAACCTGGAGGGGGGTAAAACCTCTGGGATG |
| Q5_pALFA_Lcb2_for | TTACGTCGTCGTTTGACCGAACCCATGAGTACTCCTGCAAAC |
| Q5_pALFA_Lcb2_rev | TTCCTCTTCCAACCTGGAGGGCATAATCTATTACTTTTTACTTTGAAAAC |
